# Global transcriptional reprogramming by cytomegalovirus infection suppresses MHC class II antigen presentation while enhancing migration machinery in murine dendritic cells

**DOI:** 10.1101/2025.03.31.646504

**Authors:** Rhys H. Parry, Christopher L. D. McMillan, Kimberley L. Bruce, Helen E. Farrell

**Author notes:** Correspondence: Rhys H. Parry,; Helen E. Farrell. These authors contributed equally (Rhys H. Parry, Christopher L. D. McMillan).

## Abstract

Dendritic cells (DCs) serve dual roles in cytomegalovirus infection: orchestrating antiviral immunity and acting as vehicles for viral dissemination. DC-dependent systemic spread of mouse cytomegalovirus (MCMV) is dependent on expression of a viral G protein-coupled receptor (GPCR) homolog, encoded by M33. We performed global transcriptional profiling of murine DCs infected with either wild-type mouse MCMV or a M33 mutant harbouring a single point mutation (R131Q; denoted M33NQY), which ablates constitutive G protein-dependent signalling. RNA-seq analysis revealed that MCMV induces substantial transcriptional reprogramming, with over 2,400 significantly altered genes affecting key immune and migration pathways. Wild-type MCMV infection resulted in 1,883 upregulated and 658 downregulated genes, while M33NQY infection showed comparable effects with 1,905 upregulated and 668 downregulated genes. Both viruses systematically downregulated the MHC class II antigen presentation machinery, with substantial suppression of H2 molecules, peptide-loading components (*H2-DMa/H2-DMb1*), and the class II transactivator *Ciita* (log_2_FC > -6.4). Pathway analysis revealed coordinated disruption of B cell receptor signalling, leukocyte transendothelial migration, and antigen processing and presentation. A comparison between wild-type and M33NQY-infected DCs demonstrated that while both viruses similarly impair antigen presentation, M33 signalling specifically enhances the expression of genes involved in transendothelial migration, including *Spp1* (osteopontin, log_2_FC = 0.65), cytoskeletal components (*Actg1, Actb*), adhesion molecules (*Icam1*), and matrix interaction factors (*Tnc, Plau*). Network analysis identified critical hub genes, including *B2m, Itgb1, Itgam*, and *Icam1*, as key regulatory nodes connecting antigen presentation and migration pathways. This provides molecular evidence for a sophisticated viral strategy that shields MCMV from immune detection while hijacking DC migratory machinery to facilitate dissemination, with M33 serving as a specific enhancer of migration-related pathways.

**Impact Statement:** This study provides the first comprehensive transcriptional profile of mouse cytomegalovirus (MCMV) infection in dendritic cells, revealing a sophisticated viral strategy that simultaneously suppresses antigen presentation while enhancing migration machinery. We identified that MCMV systematically downregulates the entire MHC class II pathway by targeting the master regulator *Ciita*, while the viral G protein-coupled receptor M33 specifically enhances genes involved in transendothelial migration, including *Spp1* (osteopontin). Our network analysis identified critical hub genes connecting immune evasion and migration pathways, providing molecular evidence for how MCMV exploits dendritic cells as vehicles for dissemination while evading immune detection. These findings advance our understanding of betaherpesvirus pathogenesis and highlight potential therapeutic targets to limit viral spread.

## INTRODUCTION

Dendritic cells (DCs) serve as sentinels of the immune system, initiating adaptive responses against pathogens while maintaining self-tolerance (Banchereau and Steinman, 1998; Merad et al., 2013). Strategically positioned at mucosal surfaces, DCs actively sample their environment to detect infection through pattern recognition receptors (Iwasaki, 2007). Upon pathogen encounter, DCs undergo morphological and functional changes, transitioning from a smooth, rounded state to an activated form characterised by extensive pseudopodia that facilitate their migration to draining lymph nodes (LN) (Randolph et al., 2005). This migration process is orchestrated by chemokine signalling networks that guide DCs through tissues via activation of G protein-coupled receptors (GPCRs), leading to cytoskeletal reorganisation and directional motility (Forster et al., 2008; Sozzani et al., 1998).

Within lymph nodes, migrated DCs localise to fibroblastic reticular cell niches and position themselves proximal to high endothelial venules (HEV) to facilitate interaction with incoming blood-borne T cells (Bajenoff et al., 2006; Sixt et al., 2005). In this context, mature DCs function as professional antigen-presenting cells (APCs), expressing pathogen-derived peptides via major histocompatibility complex (MHC) molecules and upregulating costimulatory signals that lower the threshold for T cell activation (Guermonprez et al., 2002; Reis e Sousa, 2006). Chemokines continue to shape DC behaviour throughout this process by engaging G protein-coupled chemokine receptors, activating downstream pathways essential for DC chemotaxis, survival, cytokine production, and differentiation (Allavena et al., 2000). Importantly, these migratory APCs have a finite lifespan in lymph nodes, disappearing rapidly from T cell areas without entering efferent lymphatics, suggesting they undergo local elimination after fulfilling their immune activation role (Ingulli et al., 1997; Kamath et al., 2002; Garg et al., 2003).

Recent investigations into the transcriptional landscape of DCs have revealed a multi-layered network of pathways that underpin the dynamic expression of DC stimulatory molecules during inflammation, particularly in response to viral challenge (Luber et al., 2010; Miller et al., 2012). These techniques offer a powerful approach to identifying key checkpoints of host immunity that can be harnessed for clinical therapies. However, several persistent viruses also use DCs as a vehicle to facilitate their dissemination, seemingly hijacking them to use as effective shields against host attack (Auffermann-Gretzinger et al., 2001). Their coexistence requires virus strategies to disrupt DC responses to extracellular inflammatory cues, which may jeopardise both virus and host cell survival. The betaherpesvirus cytomegalovirus (CMV) is among this specialised virus group (Sinclair and Reeves, 2013).

Human CMV (HCMV) is a ubiquitous pathogen that causes a largely asymptomatic infection in immunocompetent hosts (Boeckh and Geballe, 2011). In persons whose immunity is immature or immunodeficient, HCMV has the propensity to cause disseminated lytic infection and inflammation, leading to severe morbidity or death (Griffiths et al., 2015). HCMV establishes a latent infection in CD34^+^ hematopoietic myeloid progenitor cells and CD14^+^ monocytes (Reeves and Sinclair, 2013; Sinclair and Sissons, 2006). Analysis of differentially expressed genes in latently infected CD14^+^ monocytes showed marked downregulation in MHCII and *CD74* transcripts, suggestive of HCMV interference in antigen presentation and endothelial transmigration functions (Shnayder et al., 2020; Yunis et al., 2018). In contrast, the role of DCs in facilitating primary HCMV spread is less clear, as infections are asymptomatic and thus difficult to capture and track. Given the difficulty in studying early HCMV dissemination, MCMV infection in mice offers a powerful alternative for dissecting virus-DC interactions *in vivo*.

Infection of mice with mouse cytomegalovirus (MCMV) has provided a tractable approach to track virus entry and spread (Bruce et al., 2022; Krmpotic et al., 2003). Naturally acquired MCMV exploits olfaction to enter new hosts, spreading from the olfactory epithelium to draining lymph nodes via infected DC (Farrell et al., 2016). Experimental lung inoculation of MCMV results in infection of alveolar/interstitial macrophages, type II alveolar epithelial cells and DCs, with DCs remaining the principal vehicle for systemic spread (Bruce et al., 2022; Farrell et al., 2015). Remarkably, MCMV^+^ DCs are not retained in LNs. Instead, they traverse to HEV, recirculate to the blood and extravasate across vascular endothelia to numerous tissues, including the salivary glands (Farrell et al., 2017; Ma et al., 2021). Indeed, lung infection of mice with a replication-deficient (i.e., single-cycle) MCMV results in the translocation of MCMV+ DCs from the lungs to the salivary glands, which is detected seven days later. In the salivary glands, MCMV^+^ DCs transfer infection to acinar epithelial cells, where the virus is amplified, and extracellular virions are released in saliva (Farrell et al., 2017; Ma et al., 2021).

Consistent with the central role of host chemokines in DC trafficking, DC recirculation requires expression of MCMV chemokine receptors and chemokine homologs, which act in a non-redundant fashion at key stages of the DC journey (Saederup et al., 1999; Suvas and Rouse, 2006). In addition to MCMV-induced modifications in DC motility and directional decision-making, MCMV^+^ DCs exhibit other modifications, such as ablated responses to LN retention signals, diminished T cell engagement in LNs, enhanced endothelial extravasation and enhanced DC survival (Andrews et al., 2001; Mathys et al., 2003). Many MCMV genes, some of which have already been characterised in other infected cell types, will likely contribute to the altered state of DCs. However, many described examples of MCMV-induced cellular dysregulation or immune evasion occur post-transcriptionally (Loewendorf and Benedict, 2010). Little is known about the global changes in the transcriptional landscape caused by MCMV infection of DCs.

The viral GPCR (vGPCR) M33 plays a critical role in this process (Cardin et al., 2009; Davis-Poynter et al., 1997). DCs infected with MCMV lacking M33 (ΔM33) or expressing a signalling-deficient mutant (M33NQY) with a mutation in the vGPCR “DRY box motif” show markedly diminished ability to escape lymph nodes and enter the bloodstream (Farrell et al., 2017). This M33 signalling defect serves as a valuable tool to dissect M33-dependent functions throughout our study. While the phenotypic importance of M33 in DC-dependent viremia is established, the molecular mechanisms underlying this effect remain unknown. Similarly, MCMV-infected DCs exhibit altered immune functions, including reduced T cell engagement and enhanced survival, but the transcriptional basis for these changes has not been elucidated (Lemmermann et al., 2011).

In this study, we performed transcriptional profiling of murine DCs infected with either wild-type MCMV or the signalling-deficient M33NQY mutant to identify the molecular pathways underlying viral manipulation of DC biology. We hypothesised that MCMV infection would induce coordinated transcriptional changes affecting both antigen presentation and migration pathways, with M33 specifically contributing to migration-related gene expression. Our findings reveal a sophisticated viral strategy that systematically targets the MHC class II antigen presentation machinery while selectively enhancing genes involved in transendothelial migration, providing molecular insights into how MCMV exploits DCs for dissemination while evading immune detection.

Downregulation of MHC molecules at the cell surface during CMV infection has been demonstrated at the protein level in multiple settings. In murine bone-marrow–derived DCs and cDC1/cDC2 subsets, MHC-II surface abundance is reduced by ∼24–48 hpi, consistent with prior reports of class-II suppression in CMV-infected APCs. MHC-I is also targeted by CMV via multiple mechanisms (e.g., US2/US11-like pathways in HCMV; m152/m06/m04 in MCMV) that retain or degrade peptide–MHC-I complexes and limit antigen export to the surface. Here, we explicitly quantify the transcriptional repression of the class-II pathway in Mutu cDC1 and delineate how M33 signalling contributes to the migration programme while leaving class-II suppression largely intact.

## MATERIALS AND METHODS

### Cells and viruses

The mouse Mutu DC line, closely resembling conventional CD8^+^ dendritic cells but lacking detectable baseline GFP fluorescence under UV microscopy, was obtained from Michael Chopin (Monash University, Australia). Cells were cultured in Iscove’s Modified Dulbecco’s Medium (IMDM; Gibco) supplemented with 10% heat-inactivated fetal calf serum (hiFCS), 2 mM glutamine, 100 IU/mL penicillin, 100 μg/mL streptomycin, and 50 μM 2-mercaptoethanol. As with other immortalised dendritic cell models, Mutu DCs express SV40 large T antigen to support continuous growth; while this may influence some regulatory pathways, non-transformed cDC1 equivalents are not available, and Mutu cells remain a widely used model for MCMV infection studies. DCs were infected with GFP-tagged MCMV IC2 or NQY virus, which are derived from a K181 background, diluted in IMDM containing 2% hiFCS when cultures were 75% confluent. The GFP cassette is expressed from the human CMV IE1 promoter and inserted at the unique EcoRV site within the intron of MCMV m131, as previously described (Wagner et al., 1999). NQY is genetically identical to IC2 except for a point mutation in the viral GPCR M33 (R131Q; denoted M33NQY) that ablates constitutive signalling (Farrell et al., 2007; McArdle et al., 2017). IC2 replicates and spreads like untagged K181, and the NQY mutation does not impair replication in Mutu DCs. Infections (n=3) were performed at a multiplicity of 5, and all cells were GFP+ within 24 h p.i. Following a one-hour incubation with the virus, cells were washed three times with IMDM and then cultured for 48 hours in IMDM containing 2% hiFCS. Control cultures (n=3) were mock-infected and processed identically.

### RNA isolation, library preparation and next-generation sequencing

DC cultures (n = 3 per group) were harvested 2 days after infection, and RNA was isolated using the RNeasy Mini Kit according to the manufacturer’s instructions (QIAGEN). RNA integrity was assessed using an Agilent Bioanalyzer 2100 system (Agilent Technologies), and RIN values ranged from 7 to 9. Stranded mRNA sequencing libraries were sequenced using the NextSeq 150 cycle kit (Illumina Inc), generating 75-76 bp paired-end reads. The Illumina bcl2fastq 2.20.0.422 pipeline was then used to produce the sequence data. The University of Queensland Genomics Facility and the Australian Genomics Research Facility performed library preparation, sequencing, and data acquisition. Image analysis was performed on the instrument computer using HiSeq Control Software vHD 3.4.0.38 and Real-Time Analysis v2.7.7.

### Transcriptome data analysis and software

Raw, basecalled fastq files were first analysed using FastQC to assess read quality. FASTp (v0.24.0) was then used to remove adapters, low-quality reads, and reads shorter than 50 bases. For host transcriptome analysis, the remaining high-quality reads were mapped using HISAT2 (v2.2.1) to the *Mus musculus* mm10 genome (Assembly: GCA_000001635.2). Feature counts were generated with htseq-count (v0.9.1) under the following conditions: union mode, stranded: yes, minimum alignment quality: 10, feature type: exon, and ID attribute: gene_ID using the GTF file mm10.refGene.gtf (Available here: https://hgdownload.soe.ucsc.edu/goldenPath/mm10/bigZips/genes/mm10.refGene.gtf.gz). Viral transcript abundance was assessed in a separate analysis by aligning the same libraries to the Murine cytomegalovirus (strain K181; GenBank ID: AM886412) genome; no combined reference or sequential subtraction approach was used. Analyses were performed in R (v4.4.2) using RStudio (2024.12.0+467). Core packages included tidyverse (v2.0.0; Wickham et al., 2019), ggplot2 (v3.5.2; Wickham, 2009), dplyr (v1.1.4), tidyr (v1.3.1), lubridate (v1.9.4), and pheatmap (v1.0.13). Full package versions and session information are available in the GitHub repository (see sessionInfo.txt). Count files were merged for each library for individual host gene expression matrices and MCMV gene expression matrices, and differential gene expression analysis was undertaken using edgeR (v3.36.0), setting min.total.count to 10 and employing estimateDisp() with robust = TRUE, as well as glmQLFTest() with a robust fit setting. Visualisation of the fold change and FDR P-value data was achieved through volcano plots generated using EnhancedVolcano (v1.12.0). Heatmaps were generated using the pheatmap R package (v1.0.12), with default normalisation parameters where normalised expression values were scaled per gene to a z-score distribution exported and graphed using GraphPad Prism (v10.4.1). For visualisation of virus genome expression, bigWig coverage tracks were generated from merged, coordinate-sorted BAM files (Picard MergeSamFiles v3.1.1.0; Galaxy) using deepTools bamCoverage (v3.5.4; Galaxy) with a 50-bp bin size and reads-per-genomic-content (RPGC) normalisation to 1× coverage (--normalizeUsing RPGC), specifying the effective genome size of the MCMV genome reference (230,301 bp, GenBank AM886412). Tracks were written as bigWig files and viewed in Integrated Genomics Viewer (v2.7.9) with per-track log scaling.

### Gene ontology and pathway enrichment analyses and Network reconstruction

The entire differentially expressed gene set was subjected to pathway enrichment analysis using easyGSEA (Cheng et al., 2021). Multiple pathway databases were queried, including Gene Ontology (Biological Process, Cellular Component, Molecular Function), KEGG, and Reactome pathways. Gene set filtering parameters were set to include only pathways containing between 15 and 200 genes. For gene set enrichment analysis (GSEA), 1,000 permutations were performed to establish statistical significance, and pathways with P-values < 0.005 were considered significantly enriched. For network reconstruction, leading-edge DEGs from the easyGSEA analysis specifically associated with Antigen Presentation and Leukocyte Migration pathways were extracted and imported into Cytoscape (v3.10.3). Protein-protein interaction (PPI) networks were constructed using stringApp (v2.2.0) with *Mus musculus* as the reference organism. Interactions were filtered using a confidence score threshold of > 0.4, and no additional interactors were included beyond the input gene set. The individual networks for Antigen Presentation and Leukocyte Migration were then integrated using Cytoscape’s “merge networks” function with the “Union” mode to create a comprehensive interaction network. Network topology was analysed by calculating betweenness centrality values for each node using Cytoscape’s “Analyze Network” function, which quantifies the number of shortest paths passing through each node. To visualise differential expression within the network context, logFC values for each gene were incorporated as a node attribute and represented through a colour gradient.

## RESULTS

### Global transcriptomic reprogramming of murine DCs in response to MCMV infection

RNA sequencing of murine dendritic cells revealed extensive transcriptional reprogramming following infection with either wild-type MCMV (IC2) or the signaling-deficient M33NQY mutant. Mock-infected samples yielded an average of 16.9 million reads per library with high mapping efficiency to the mouse genome (97.26%). In contrast, both infected conditions showed a reduction in host reads due to viral transcription: IC2-infected samples contained 14.9 million reads (75.81% mapping to mouse genome, 18.46% to MCMV genome), while M33NQY-infected samples showed a similar distribution (14.1 million reads; 76.90% mouse, 17.15% MCMV) (Fig. 1A, Table S1). Despite these differences in raw mapping percentages, edgeR normalization effectively adjusted for these technical variations (Fig. 1B).

**Figure 1.**
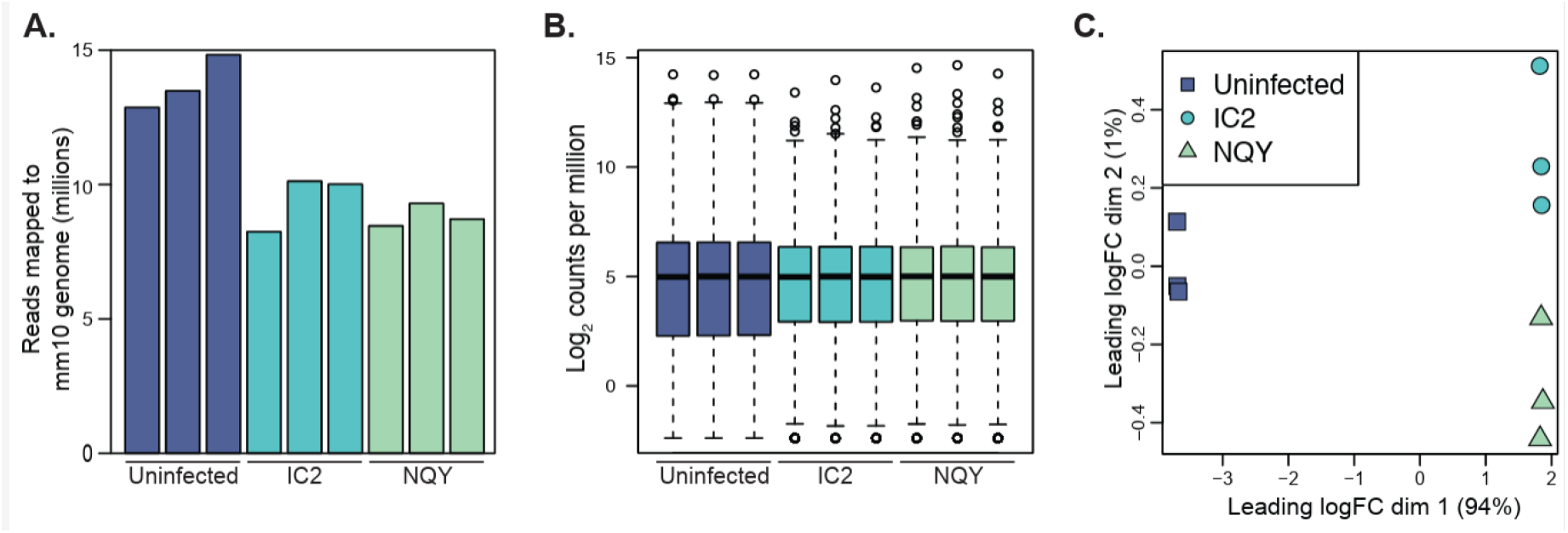
Transcriptional profiling of MCMV-infected dendritic cells. (A) Summary of reads mapped to genes in the mm10 genome from dendritic cells (DCs) mock-infected or infected with either wild-type MCMV (IC2) or the M33NQY mutant (n=3 per group). (B) edgeR-normalised library size across samples. (C) Multidimensional scaling plot of the normalised libraries showing the relationships between biological replicates of each condition.

Multidimensional scaling analysis (Fig. 1C) demonstrated clear separation between infected and uninfected conditions along dimension 1 (capturing 94% of total variation), with dimension 2 (capturing 1% of variation) distinguishing between IC2 and M33NQY infections. Biological replicates within each condition clustered tightly together, confirming high reproducibility.

Our analysis of MCMV-infected versus mock-infected dendritic cells revealed massive transcriptional reprogramming, with 2,541 genes showing significant differential expression in wild-type (IC2) infection using stringent criteria (|log_2_FC| > 2 and FDR < 0.05) (Fig. 2A; Table S2). Similarly, the M33NQY mutant virus induced substantial transcriptional changes, with 2,573 genes significantly altered (Fig. 2A, Table S3). Wild-type MCMV infection resulted in 1,883 significantly upregulated and 658 significantly downregulated genes, while M33NQY infection showed comparable effects with 1,905 upregulated and 668 downregulated genes (Fig. 2A).

**Figure 2.**
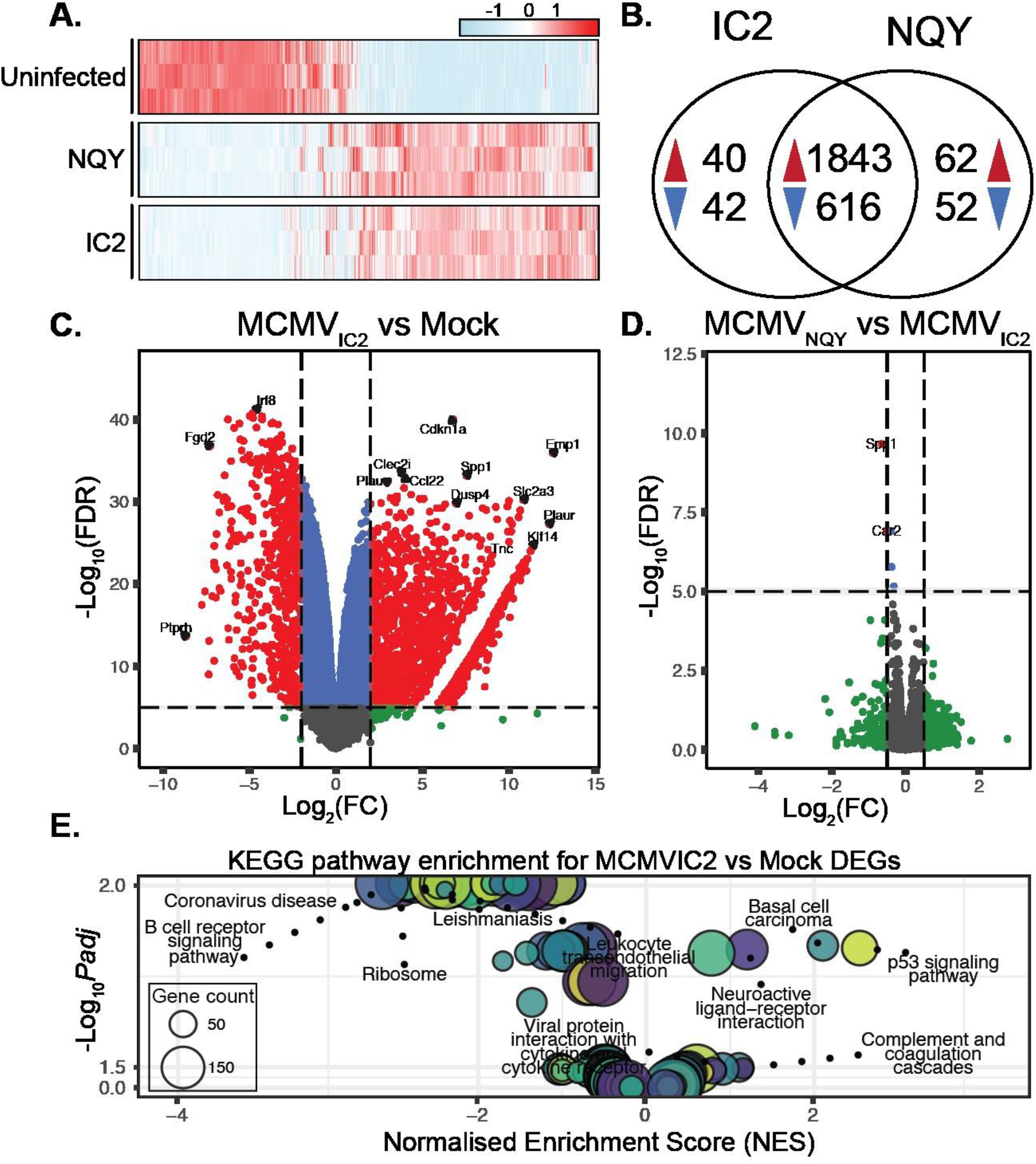
Differential gene expression in MCMV-infected dendritic cells. (A) Heatmap of normalised expression (Z-scores) for differentially expressed genes, scaled per gene and hierarchically clustered (Euclidean distance, complete linkage) by gene only; sample order is preserved. (B) Venn diagram showing overlap of differentially expressed genes (DEGs) in dendritic cells infected with wild-type MCMV (IC2) or M33NQY mutant compared with mock controls. Upregulated genes are indicated in red (▴) and downregulated genes in blue (▾). Numbers outside the overlap represent DEGs unique to each infection; numbers within the overlap represent DEGs shared between IC2 and NQY infections. (C) Volcano plot of differentially expressed genes in wild-type (IC2) MCMV-infected DCs compared with mock controls. For visualisation, a stringent cutoff of FDR ≤ 10^−5^ is shown to highlight the most robust hits (overall significance for differential expression is defined as FDR < 0.05). Red: FDR ≤ 10^−5^ and |log_2_FC| > 2; blue: FDR ≤ 10^−5^ but |log_2_FC| ≤ 2; green: FDR > 10^−5^ but |log_2_FC| > 2; grey: not significant. Selected immune-related genes are labelled. Only genes with a total count > 10 were included (n = 12,805). Y-axis shows – log_10_(FDR). (D) Volcano plot depicting differential gene expression between DCs infected with wild-type MCMV (IC2) and the M33NQY mutant. (E) Functional enrichment analysis of MCMVIC2 vs Mock differentially expressed genes showing significantly enriched KEGG pathways. Bubble size indicates the number of differentially expressed genes contributing to each pathway (gene count), the y-axis shows statistical significance (–log_10_(FDR)), and the x-axis shows normalised enrichment score (NES). The top five positively and negatively enriched pathways are labelled. Bubble colour is arbitrary and used only to visually separate points.

Comparative analysis revealed remarkably high concordance between wild-type and M33NQY mutant infections, with the vast majority of transcriptional changes conserved between the two viruses (Fig. 2B). Of the differentially expressed genes, 2,459 genes (96.8%) showed significant alterations in both infections, including 1,843 genes that were upregulated and 616 genes that were downregulated in both conditions. Only a small subset of genes displayed strain-specific responses: 40 genes were uniquely upregulated and 42 genes uniquely downregulated in IC2 infection. In comparison, 62 genes were uniquely upregulated and 52 genes uniquely downregulated in M33NQY infection (Fig. 2B). This high degree of overlap (>93% for downregulated genes and >96% for upregulated genes) indicates that the M33 NQY mutation does not substantially alter the core transcriptional response of dendritic cells to MCMV infection.

Both viruses induced similar patterns of extreme gene expression changes (IC2 versus mock Fig. 2C), with top upregulated genes in both infections including *Emp1* (log_2_FC = 12.58 in IC2, 12.49 in NQY), *Plaur* (log_2_FC = 12.44 in IC2, 12.19 in NQY), and *Klf14* (log_2_FC = 11.35 in IC2, 11.37 in NQY). Both viruses strongly suppressed immune-related genes, including *Ptprh* (log_2_FC = -8.66 in both strains), *Fgd2* (log_2_FC = -7.18 in IC2, -7.38 in NQY), and *Clec9a* (log_2_FC = -7.04 in IC2, -7.24 in NQY). The downregulation of *Clec9a* is particularly noteworthy as this C-type lectin receptor is specifically expressed by conventional dendritic cells and is crucial for cross-presentation of necrotic cell-associated antigens to CD8+ T cells (Sancho et al., 2009; Zelenay et al., 2012; Zhang et al., 2012).

The comparison between wild-type (IC2) and M33NQY mutant MCMV-infected dendritic cells revealed far more subtle transcriptional differences. Using less stringent criteria (|log_2_FC| > 0.5 and FDR < 0.05), we identified only 35 significantly differentially expressed genes (Fig. 2D; Table S4). Of these, 22 genes were upregulated and 13 genes were downregulated in IC2 compared to NQY-infected cells. The most differentially expressed gene was the lincRNA *2310001H17Rik*, which is currently uncharacterised (log_2_FC = 2.18), followed by several immune-related genes, including *Ifit1bl1* (log_2_FC = 1.53), *Apol7c* (log_2_FC = 1.37), and the complement component *C3* (log_2_FC = 1.34). Notably, *Nos2* (inducible nitric oxide synthase) was downregulated in IC2 compared to NQY-infected cells (log_2_FC = -1.16), potentially indicating differences in inflammatory response regulation between the two viral strains.

### Pathway analysis reveals coordinated targeting of antigen presentation and cell migration processes

KEGG pathway enrichment analysis identified significant (padj < 0.01) downregulation in multiple immune and migration-related pathways in both wild-type and M33NQY mutant infections (Fig. 2E; Tables S5, S6). For wild-type MCMV, the most downregulated pathways included Coronavirus disease (NES = -2.50), B cell receptor signalling pathway (NES = -2.43), Ribosome (NES = -2.40), and Leukocyte transendothelial migration (NES = -2.36). In M33NQY-infected cells, Fc gamma R-mediated phagocytosis ranked first (NES = -2.47), followed by Coronavirus disease (NES = -2.42), Leukocyte transendothelial migration (NES = -2.41), and B cell receptor signalling pathway (NES = -2.40). Antigen processing and presentation were also significantly downregulated in both conditions (NES = -2.26 in IC2, rank #7; NES = -2.27 in NQY, rank #6).

We also observed significant upregulation (padj < 0.05) of several pathways. The p53 signalling pathway showed the highest positive enrichment in both infections (NES = 2.22 in IC2 and NES = 2.26 in NQY), suggesting the activation of stress response mechanisms. Other upregulated pathways included Complement and coagulation cascades (NES = 1.70 in IC2; NES = 1.72 in NQY), Viral protein interaction with cytokine and cytokine receptor (NES = 1.68 in IC2; NES = 1.67 in NQY), and Cytokine-cytokine receptor interaction (NES = 1.58 in IC2; NES = 1.61 in NQY).

Gene Ontology biological process analysis further supported these findings, with significant downregulation in terms related to antigen processing and presentation (GO:0019882, NES = - 2.63 in IC2, -2.62 in NQY), T cell activation (GO:0042110, NES = -2.41 in IC2, -2.39 in NQY), and leukocyte migration (GO:0050900, NES = -2.29 in IC2, -2.26 in NQY). Broader immune system processes were also suppressed in both infections, including leukocyte activation (NES = -2.35 in IC2 and -2.33 in NQY) and regulation of immune system processes (NES = -2.24 in IC2 and -2.30 in NQY). Both viruses showed significant upregulation of biological processes related to DNA metabolism (NES = 2.68 in IC2, 2.73 in NQY), DNA replication (NES = 2.65 in IC2, 2.68 in NQY), and cell cycle processes (NES = 2.46 in IC2, 2.52 in NQY), possibly reflecting viral hijacking of host cellular machinery to support MCMV genome replication.

### Comparing wild-type MCMV and M33NQY mutant MCMV transcriptional effects

KEGG pathway analysis comparing WT MCMV (IC2) versus the M33NQY mutant revealed a selective pattern of enrichment with a distinct emphasis on cell migration and cytoskeletal organisation pathways (Table S6). Leukocyte transendothelial migration showed the strongest positive enrichment (NES = 2.11, padj < 0.05), followed by Fc gamma R-mediated phagocytosis (NES = 2.09, padj < 0.05), Focal adhesion (NES = 2.05, padj < 0.05), and Regulation of actin cytoskeleton (NES = 1.97, padj < 0.05). These findings align with previous work demonstrating M33’s critical role in DC migration and dissemination (Farrell et al., 2017; Ma et al., 2022).

While the comparison between the wild-type and M33NQY mutant revealed relatively subtle transcriptional differences (only 35 genes with |log_2_FC| > 0.5 and FDR < 0.05, Fig. 2D), we identified significant changes in key molecules related to migration. The most statistically significant difference was observed in *Spp1* (osteopontin, log_2_FC = 0.65, FDR < 1E-9), a secreted glycoprotein critical for leukocyte migration and DC maturation (Renkl et al., 2005). Other significantly upregulated migration-related genes in wild-type MCMV included cytoskeletal components (*Actg1*, log_2_FC = 0.26, FDR < 0.0001; *Actb*, log_2_FC = 0.19, FDR < 0.002), cell adhesion and matrix interaction factors (*Tnc*, log_2_FC = 0.56, FDR < 0.0001; *Plau*, log_2_FC = 0.39, FDR < 1E-7), cytoskeletal regulators (*Gsn*, log_2_FC = 0.32, FDR < 0.0001; *Rac2*, log_2_FC = 0.18, FDR < 0.01), and adhesion proteins (*Icam1*, log_2_FC = 0.17, FDR < 0.03). Notably, *Car2* (log_2_FC = 0.52, FDR < 1E-7) was also significantly upregulated, suggesting potential involvement of pH regulation mechanisms. The lincRNA *2310001H17Rik* (log_2_FC = 2.18, FDR < 0.03) showed the largest fold change, followed by the interferon-inducible gene *Ifit1bl1* (log_2_FC = 1.53, FDR < 0.01) and complement component *C3* (log_2_FC = 1.34, FDR < 0.03), suggesting additional regulatory mechanisms. Similar upregulation of migration-associated genes has been observed in HCMV-infected endothelial cells expressing the homologous US28 vGPCR (Maussang et al., 2009; Vomaske et al., 2009). While some highlighted genes (e.g., Actg1, Icam1) show more modest fold changes than our formal significance threshold, they are included to illustrate coherent pathway-level changes; overall migration pathway enrichment remains strongly supported by the analysis (Table S6).

Downregulated pathways in the wild-type versus M33NQY comparison were primarily related to ribosomal functions and metabolic processes, with Ribosome showing the strongest negative enrichment (NES = -2.29, padj < 0.05). Antigen processing and presentation pathways showed no significant differences between wild-type and M33NQY infections (NES = -1.25, padj > 0.25), suggesting that M33 signalling selectively enhances migration without affecting MCMV’s immune evasion mechanisms.

### MCMV systematically targets the MHC class II antigen presentation pathway

Our transcriptomic analysis revealed coordinated downregulation across multiple components of the MHC class II antigen presentation pathway in MCMV-infected dendritic cells (Fig. 3A), consistent with previous observations of MHC class II suppression by cytomegaloviruses (Lee et al., 2011; Tomazin et al., 1999; Yunis et al., 2018). The core MHC class II molecules showed profound downregulation in both wild-type and M33NQY mutant infections, with *H2-Aa* (log_2_FC = -4.77 in IC2; -4.82 in NQY), H2-Ab1 (log_2_FC = -4.99 in both strains), and *H2-Eb1* (log_2_FC = - 4.32 in IC2; -4.20 in NQY) all showing significant reduction. Peptide loading machinery components were similarly suppressed, including *H2-DMa* (log_2_FC = -5.55 in IC2; -5.36 in NQY), *H2-DMb1* (log_2_FC = -4.54 in IC2; -4.44 in NQY), H2-DMb2 (log_2_FC = -7.58 in IC2; -6.22 in NQY), and *H2-Ob* (log_2_FC = -2.90 in IC2; -2.51 in NQY).

**Figure 3.**
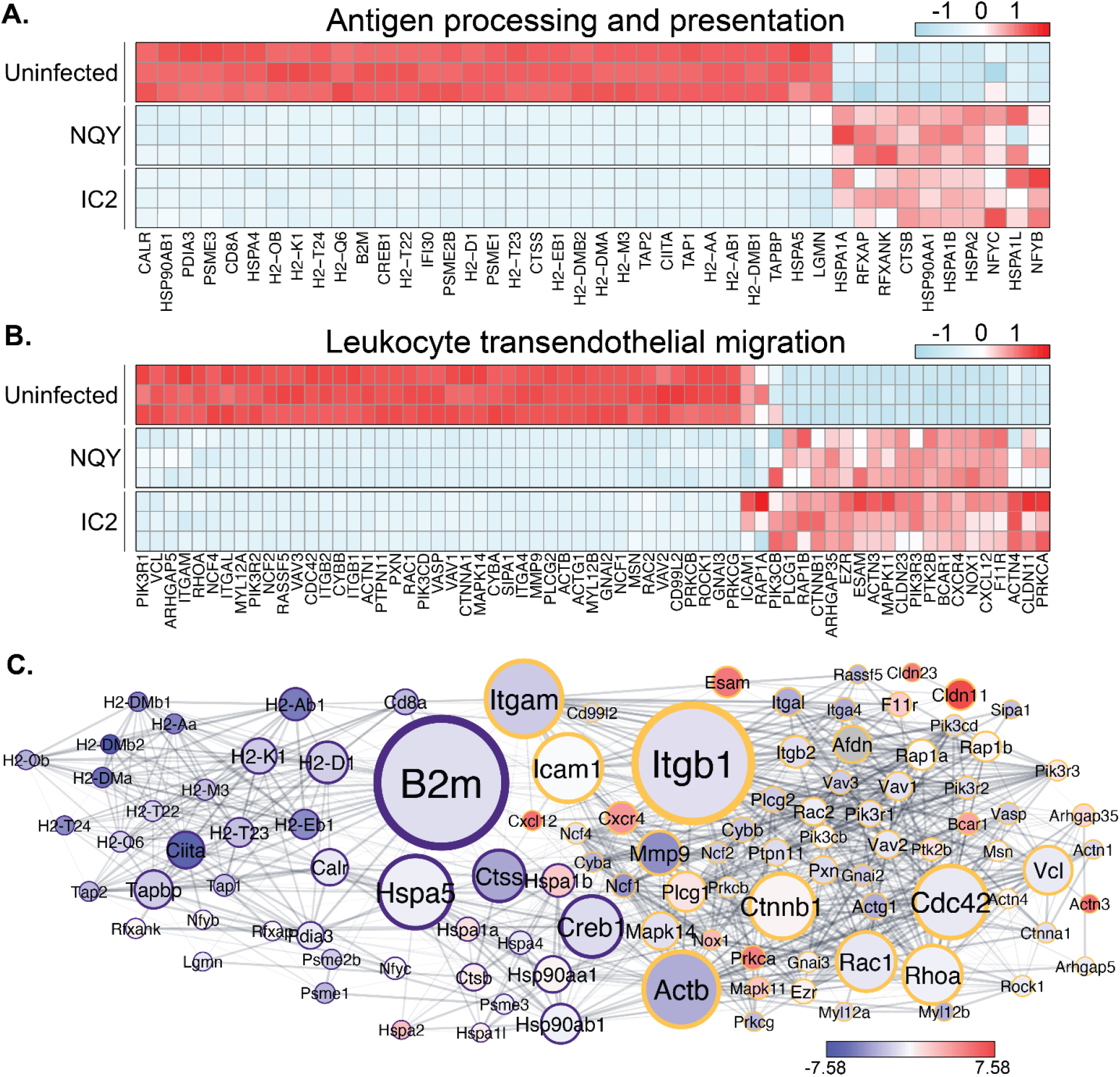
MCMV infection downregulates MHC class II antigen presentation machinery and leukocyte migration in dendritic cells. Heatmap of leading-edge genes from the (A) KEGG Antigen Processing and Presentation pathway or (B) KEGG Leukocyte Transendothelial Migration pathway across experimental conditions. Values are Z-scores of normalised expression scaled per gene and hierarchically clustered (Euclidean distance, complete linkage) by gene only; sample order is preserved. (C) Network analysis of differentially expressed genes involved in antigen presentation and leukocyte migration pathways. Purple rings: genes in the antigen presentation pathway; Yellow rings: genes in the leukocyte migration pathway. Node size represents betweenness centrality, node colour indicates log_2_FC, and edge thickness corresponds to STRING interaction confidence score.

Most notably, the Class II Transactivator (*Ciita*), the master regulator of MHC class II expression, showed substantial downregulation (log_2_FC = -6.45 in IC2; -7.03 in NQY), as did the invariant chain *Cd74* (log_2_FC = -5.49 in IC2; -6.02 in NQY), which plays multiple roles in MHC class II trafficking and peptide loading. This targeting of CIITA is reminiscent of the strategy employed by HCMV, which inhibits IFN-γ-induced upregulation of CIITA in infected cells (Le Roy et al., 1999; Lee et al., 2011).Transcriptional regulators of MHC class II expression showed varied responses, with *Rfx5* downregulated (log_2_FC = -1.26 in IC2; -1.30 in NQY), while *Rfxank* (log_2_FC = 0.28 in IC2; 0.36 in NQY) and *Rfxap* (log_2_FC = 0.17 in IC2; 0.24 in NQY) showed modest upregulation.

The overall pattern of comprehensive MHC class II pathway downregulation was conserved between the two viral strains, suggesting that this immune evasion strategy operates independently of M33 signalling.

### MCMV modulates leukocyte migration, and MCMV NQY infection attenuates migration-associated gene expression

Our GSEA identified the KEGG Leukocyte Transendothelial Migration pathway as significantly altered in MCMV-infected DCs. Heatmap visualisation of the leading-edge genes from this pathway (Fig. 3B) demonstrates changed expression in both IC2 and NQY-infected cells, although with distinct patterns between the two viral strains. Integrins exhibited mixed regulation patterns, with Itgb1 being upregulated (log_2_FC = 0.86) and Itgb2 being significantly downregulated (log_2_FC = -1.43), suggesting a selective modulation of specific adhesion pathways.

Cytoskeletal regulators, critical for DC migration, showed substantial alterations. The actin cytoskeleton components *Actb* (log_2_FC = 0.92) and *Actg1* (log_2_FC = 0.88) were upregulated, while key regulators of cytoskeletal dynamics, including *Rac1* (log_2_FC = 0.76) and *Rac2* (log_2_FC = 0.68), also showed increased expression. These Rho GTPases serve as molecular switches that coordinate the cytoskeletal remodelling necessary for cell migration.

Signalling molecules governing directional migration exhibited altered expression patterns, with PI3K components showing mixed regulation (*Pik3cd*, log_2_FC = 1.14; *Pik3r1*, log_2_FC = -0.83; *Pik3r2*, altered in both infections). The ERM (ezrin-radixin-moesin) family proteins, which link plasma membrane proteins to the actin cytoskeleton, were also modulated, with *Msn* and *Ezr* showing differential expression between IC2 and NQY infections.

Many migration-related genes exhibit stronger expression in IC2-infected cells than in NQY-infected cells (Fig. 3B), suggesting that functional M33 signalling enhances the expression of specific migration-related genes, consistent with the known role of M33 in promoting DC migration during MCMV infection.

### Integrated network analysis of antigen presentation and leukocyte migration

Network analysis of the leading-edge genes from the KEGG Antigen Processing and Presentation and Leukocyte Transendothelial Migration pathways revealed key regulatory nodes connecting these components (Fig. 3C). The network visualisation demonstrated extensive interconnectivity between antigen presentation machinery components and leukocyte migration regulators, with several hub genes exhibiting high betweenness centrality values.

*B2m* (beta-2-microglobulin) emerged as the most critical hub gene in the network, displaying the highest betweenness centrality. While primarily known as an essential component of MHC class I molecules, B2m showed significant downregulation in MCMV-infected DCs, potentially affecting both antigen presentation and cell adhesion processes. Three additional major hub genes, *Itgb1* (integrin subunit beta 1), *Itgam* (integrin alpha M, CD11b), and *Icam1* (intercellular adhesion molecule 1), formed a highly interconnected cluster at the interface of immune recognition and cellular migration pathways. *Itgb1*, which mediates cell-cell and cell-matrix interactions essential for leukocyte trafficking, showed moderate upregulation in MCMV infection. Similarly, *Icam1*, a critical endothelial adhesion molecule that facilitates leukocyte transendothelial migration, displayed increased expression, particularly in wild-type MCMV infection compared to M33NQY infection.

The network analysis also identified significant secondary hubs, including Rho GTPases (*Rac1, Rhoa, Cdc42*) and cytoskeletal components (*Actb, Actg1*), which coordinate the cellular machinery required for MHC trafficking and DC motility. These interconnected nodes represent potential convergence points where MCMV can simultaneously modulate antigen presentation and migration capabilities of infected dendritic cells.

### MCMV gene expression patterns in infected dendritic cells

Our RNA-seq analysis of MCMV-infected dendritic cells at 48 hours post-infection revealed extensive expression of MCMV genes across all temporal classes of viral genes, indicating productive infection and replication (Fig. 4A). After MCMV gene count and normalisation (Fig. 4B), individual transcript abundance of MCMV genes included high expression of immune evasion genes (m04, m06, m152, m144, m157), chemokine and cytokine modulators (m131/129, m133, m145), and tegument proteins including pp71 (M82). Multidimensional scaling analysis demonstrated clear clustering of biological replicates for both virus strains (Fig. 4C).

**Figure 4.**
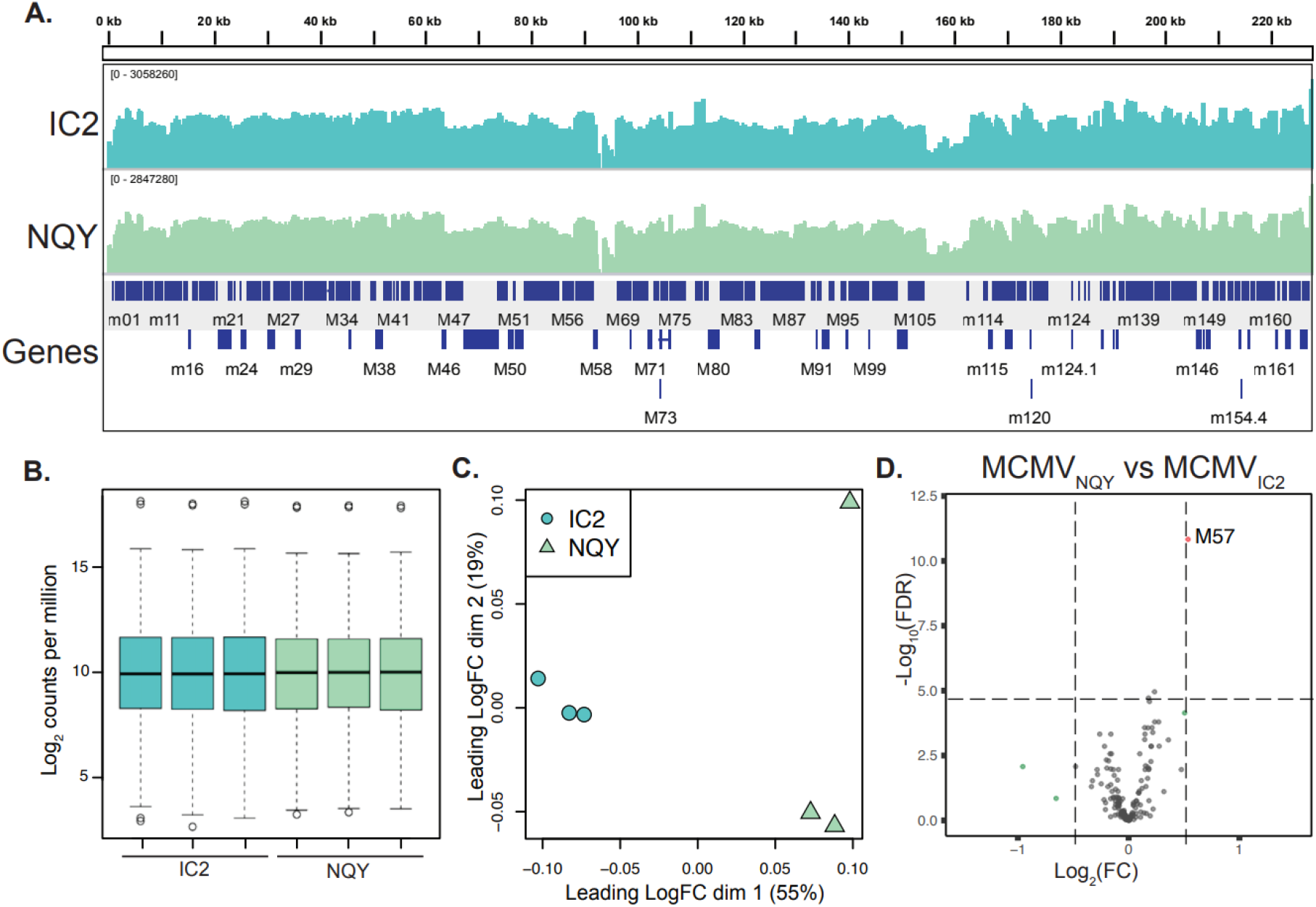
Transcriptional expression of the MCMV genome. (A) Genome browser view of RNA-seq coverage across the MCMV K181 genome (GenBank AM886412) at 48 hpi for wild-type IC2 (top) and M33NQY (middle). Tracks are merged from three biological replicates per condition, displayed as bigWig coverage in IGV with per-track log scale and auto-scaling. Annotated MCMV genes are shown below. (B) Boxplot of normalised MCMV genes per library. (C) Multidimensional scaling plot showing relationships between biological replicates of IC2-infected and M33NQY-infected DCs. (D) Volcano plot of differentially expressed MCMV genes between wild-type (IC2) and M33NQY-infected DCs. For visualisation, a stringent cutoff of FDR ≤ 10^−5^ is shown to highlight the most robust hits (overall significance for differential expression is defined as FDR < 0.05). Red: FDR ≤ 10^−5^ and |log_2_FC| > 0.5; blue: FDR ≤ 10^−5^ but |log_2_FC| ≤ 0.5; green: FDR > 10^−5^ but |log_2_FC| > 0.5; grey: not significant. n = 159 MCMV genes.

There was no difference in the expression of M33 in the M33NQY amino-acid substitution mutant, and transcript abundance was comparable between strains (mean TMM-normalised counts, IC2: 2,076 ± 41 vs NQY: 2,219 ± 65; n=3 per group) and was not called differentially expressed in our analysis. This suggests that the previously reported migratory phenotype of M33NQY-infected DCs results from a signalling defect rather than altered viral gene expression. Comparison of viral gene expression profiles revealed largely similar patterns across the viral genome (Fig. 4D). Only the M57 gene, which encodes the viral major capsid protein, showed modestly higher expression in the M33NQY mutant compared to the wild-type virus (log_2_FC = 0.58, FDR < 0.05).

This minimal difference in viral gene expression supports our conclusion that the observed phenotypic and host transcriptional differences between wild-type and M33NQY infections stem specifically from the M33 signalling defect rather than from alterations in the viral transcriptional program.

## Discussion

Our global transcriptional analysis of MCMV-infected dendritic cells reveals a sophisticated viral strategy that simultaneously impairs antigen presentation while selectively enhancing migration capabilities, facilitating viral dissemination and evading immune detection. Three key findings emerge from our data: (1) MCMV systematically downregulates the entire MHC class II antigen presentation machinery, (2) M33 signalling specifically enhances the expression of genes involved in transendothelial migration, and (3) these dual mechanisms operate largely independently, with M33 having minimal impact on MHC class II suppression.

The coordinated downregulation of multiple MHC class II pathway components represents a comprehensive immune evasion strategy that targets a central mechanism of adaptive immunity. The particularly strong suppression of *Ciita* (log_2_FC > -6.4), the master transcriptional regulator of MHC class II genes, suggests this represents a strategic viral target rather than a nonspecific effect of infection, consistent with HCMV-infected CD34(+) dendritic cells (Lee et al., 2011). By targeting CIITA, MCMV can simultaneously suppress multiple components of the MHC class II machinery through a single regulatory node. This finding extends previous observations in HCMV-infected monocytes, where MHC II and *Cd74* transcripts were diminished, demonstrating a more comprehensive suppression in actively infected DCs across the entire antigen presentation pathway. This represents a significant evolutionary adaptation, allowing MCMV to shield itself from CD4+ T cell recognition precisely within the specialised cells for initiating these responses.

Our comparative analysis between wild-type and M33NQY MCMV revealed the specific contribution of M33 signalling to the enhancement of DC migration capabilities. While the overall transcriptional differences were subtle (only 35 genes with |log_2_FC| > 0.5 and FDR < 0.05), the most statistically significant changes occurred in genes critical for cell migration. Most notably, *Spp1* (osteopontin, log_2_FC = 0.65, FDR < 1E-9), a secreted glycoprotein essential for leukocyte migration and DC maturation, showed the highest statistical significance of all differentially expressed genes. Given its role in both DC maturation and motility, the marked M33-dependent upregulation of *Spp1* likely contributes directly to the enhanced tissue dissemination observed *in vivo*. The M33-dependent upregulation of cytoskeletal components (*Actg1*, log_2_FC = 0.26, FDR < 0.0001; *Actb*, log_2_FC = 0.19, FDR < 0.002), cell adhesion and matrix interaction factors (*Tnc*, log_2_FC = 0.56, FDR < 0.0001; *Plau*, log_2_FC = 0.39, FDR < 1E-7), and adhesion molecules (*Icam1*, log_2_FC = 0.17, FDR < 0.03) provides the molecular basis for previously observed defects in lymph node escape by M33NQY-infected DCs (Ma et al., 2022). The remarkable specificity of M33’s effects, which enhances migration pathways without substantially affecting viral gene expression or immune evasion mechanisms, indicates that this vGPCR has evolved to modulate discrete host cellular functions that benefit viral dissemination.

Our network analysis revealed critical regulatory nodes bridging antigen presentation and migration pathways, with B2m emerging as the most central hub gene, a position that largely reflects its extensive connectivity to MHC molecules within the STRING/KEGG network and should be interpreted as topological rather than implying a novel direct functional link to class II components. The prominent position of *B2m* in our network suggests MCMV’s modulation of this MHC class I component may have broader implications beyond CD8+ T cell evasion, potentially affecting cell adhesion and migration processes (Farrell et al., 2019). Functional studies have demonstrated that *B2m* can act beyond its classical role in MHC class I presentation, directly regulating cell adhesion and migration through interaction with integrin signalling pathways (Huang et al., 2008). This broader signalling role of *B2m* may explain its position as a central hub connecting antigen presentation and migration networks in our analysis

The identification of *Itgb1, Itgam*, and *Icam1* as major secondary hubs demonstrates the interconnected nature of immune recognition and cellular migration, providing molecular targets through which MCMV can efficiently coordinate its dual-modulation strategy. The cytoskeletal regulatory components, including *Rac1, Rhoa*, and *Cdc42*, further highlight convergence points where viral manipulation can simultaneously impact both antigen presentation and migration capabilities. Notably, the human cytomegalovirus (HCMV) vGPCR encoded by US28, which possesses a signalling profile similar to M33 (Waldhoer et al., 2002), also converges with the RhoA GTPase signal transduction pathway (Melnychuk et al., 2004) important for cellular migration, suggesting conserved functionality.

Our findings have broader implications for understanding betaherpesvirus pathogenesis and host-pathogen interactions. The identification of M33 as a specific enhancer of migration-related pathways, rather than a general regulator of viral gene expression, highlights how viral proteins can evolve specialised functions that appropriate normal cellular processes (White et al., 2022). The systematic targeting of the MHC class II pathway, particularly through *Ciita* suppression, reveals a strategic viral approach to immune evasion that maximises effectiveness while minimising the genetic investment required.

The strong association of Spp1 with M33-dependent migration enhancement presents a particularly intriguing target for potential intervention. As the most significantly differentially expressed gene between wild-type and M33NQY-infected cells, osteopontin likely plays a central role in facilitating the enhanced migratory capabilities of MCMV-infected DCs. The importance of this glycoprotein in both migration and DC maturation suggests it could serve as a valuable biomarker or therapeutic target specific to the viral dissemination phase. Targeting *Spp1* or its downstream effectors might provide a strategy to limit viral spread while minimising disruption to normal immune functions (Icer and Gezmen-Karadag, 2018).

This transcriptomic analysis provides a molecular roadmap of how MCMV manipulates dendritic cell biology to facilitate dissemination. By identifying specific pathways and regulatory nodes targeted by viral mechanisms, our findings highlight potential intervention points for preventing viral dissemination and enhancing immune recognition of infected cells. The central role of *B2m, Itgb1, Itgam*, and *Icam1* in connecting immune and migration networks suggests these molecules might represent valuable targets for therapeutic intervention. Future studies should focus on elucidating the specific viral factors responsible for *Ciita* suppression and determining how M33 signalling selectively enhances migration-related gene expression through these key regulatory nodes. While our findings define robust transcriptional reprogramming in response to MCMV infection in DCs, we acknowledge that transcript levels do not always directly translate to protein abundance. Future work integrating proteomic or targeted protein-level validation would strengthen understanding of how these transcriptional changes manifest functionally.

## Supporting information

Table S2: S2_Mockvs_IC2.csv Differentially expressed genes in wild-type (IC2) MCMV infected cells compared to Mock.

Table S3: S3_Mockvs_NQY.csv Differentially expressed genes in M33NQY MCMV infection compared to Mock.

Table S4: S4_IC2vsNQY_DEGs.csv Differentially expressed genes between MCMV-IC2-infected versus MCMV-NQY-infected cells

Table S5: S5_Mockvs_IC2_KEGG-RA-BP-CC-MF_IC2_vs_Mock.csv easyGSEA results of wild-type (IC2) MCMV infected cells compared to Mock.

Table S6: S6_Mockvs_NQY_KEGG-RA-BP-CC-MF_NQY_vs_Mock.csv easyGSEA results in M33NQY MCMV infected cells compared to Mock.

Table S7: S7_IC2vsNQY_DEGs_KEGG-RA-BP-CC-MF_IC2_verses_NQY.csv easyGSEA results in MCMV-IC2-infected versus MCMV-NQY-infected cells.

## Data summary

The RNA sequencing data generated for this study have been deposited in the NCBI Sequence Read Archive (SRA) under accession number PRJNA1071051. Count data for mouse and MCMV genes session environment data and R scripts are available via GitHub at github.com/rhparry/MCMV_transcriptome/

## Acknowledgements

This work utilised the Australian Galaxy service (https://usegalaxy.org.au/). We acknowledge resources prepared by Dr. Andrii Slonchak for help with Cytoscape analysis, visualisation and interpretation.

## Author contributions

H.E.F conceived the study. K.B. and H.E.F collected the primary samples. H.E.F secured funding. R.H.P and C.L.D.M performed the analyses. R.H.P and C.L.D.M provided visualisation. R.H.P., C.LD.M, and H.E.F drafted the manuscript. H.E.F provided resources. All authors reviewed and edited the manuscript.

## SUPPLEMENTARY INFORMATION

**Supplementary Table 1:**
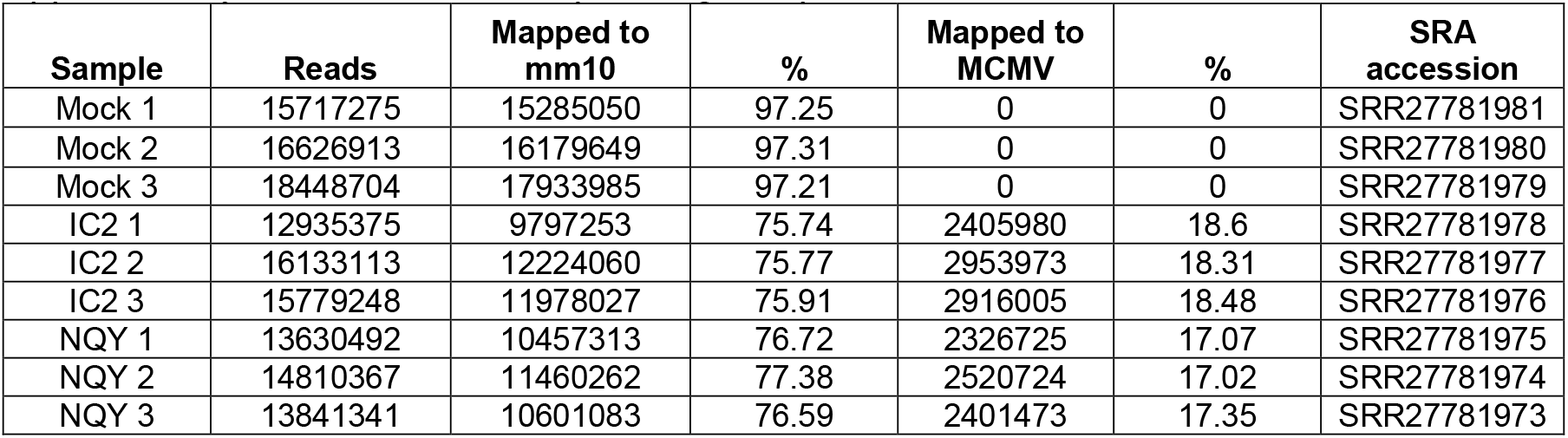
mRNA sequencing sample metrics

- Table S2: *S2_Mockvs_IC2*.*csv* Differentially expressed genes in wild-type (IC2) MCMV infected cells compared to Mock.
- Table S3: *S3_Mockvs_NQY*.*csv* Differentially expressed genes in M33NQY MCMV infection compared to Mock.
- Table S4: *S4_IC2vsNQY_DEGs*.*csv* Differentially expressed genes between MCMV-IC2-infected versus MCMV-NQY-infected cells
- Table S5: *S5_Mockvs_IC2_KEGG-RA-BP-CC-MF_IC2_vs_Mock*.*csv* easyGSEA results of wild-type (IC2) MCMV infected cells compared to Mock.
- Table S6: *S6_Mockvs_NQY_KEGG-RA-BP-CC-MF_NQY_vs_Mock*.*csv* easyGSEA results in M33NQY MCMV infected cells compared to Mock.
- Table S7: *S7_IC2vsNQY_DEGs_KEGG-RA-BP-CC-MF_IC2_verses_NQY*.*csv* easyGSEA results in MCMV-IC2-infected versus MCMV-NQY-infected cells.

